# Mendelian Randomization with Instrumental Variable Synthesis (IVY)

**DOI:** 10.1101/657775

**Authors:** Zhaobin Kuang, Aldo Cordova-Palomera, Fred Sala, Sen Wu, Jared Dunnmon, Chris Re, James Priest

**Affiliations:** Stanford University

## Abstract

Mendelian Randomization (MR) is an important causal inference method primarily used in biomedical research. This work applies contemporary techniques in machine learning to improve the robustness and power of traditional MR tools. By denoising and combining candidate genetic variants through techniques from unsupervised probabilistic graphical models, an influential latent instrumental variable is constructed for causal effect estimation. We present results on identifying relationships between biomarkers and the occurrence of coronary artery disease using individual-level real-world data from UK-BioBank via the proposed method. The approach, termed Instrumental Variable sYnthesis (IVY) is proposed as a complement to current methods, and is able to improve results based on allele scoring, particularly at moderate sample sizes.

## 1. INTRODUCTION

The goal of causal inference is to estimate the true causal relationship between a risk factor and an outcome from observational data in the absence of a controlled experiment (Judea Pearl, Madelyn Glymour, 2016). Using genetic data derived from large-scale genome-wide association studies (GWAS), Mendelian Randomization (MR) is an important method for causal inference in biomedical research. Causal relationships in observational data may be obscured by confounders, which are intermediate phenomena associated with both the risk factor and the outcome. Under the assumption that genetic variant A (inherited randomly from a parent) is associated with a risk factor, but does not confer risk directly for the outcome of interest or a confounder, the presence of confounders in observational data can be overcome by using genetic variant A in MR to infer the true causal relationship. A genetic variant conferring risk for a trait meeting these criteria is called an instrumental variable (IV)—a variable which influences a risk factor independent of the confounders, and subsequently influences the outcome only through the risk factor (Angrist, Imbens, & Rubin, 1996). Mendelian Randomization is among the most popular applications of instrumental variable methodology to real-world problems in biomedical science (Palmer et al., 2012).

The development of MR methods has been centered around publicly available genetic data. In the era of GWAS most studies release summary statistics, which report β-coefficients relating individual genetic variants to one or more related outcomes or traits. Using summary statistics, there is a proliferation of methods to test the underlying assumptions of independence of genetic variants from confounders and outcomes, and to perform MR (Burgess, Small, & Thompson, 2017). The sample size required for well-powered MR is proportional to the amount of variance explained (r2) by both the genetic instrumental variable on the risk factor, and the strength of the relationship of between the risk factor to the outcome itself. With an r2 typically less than 0.01 derived from GWAS summary statistics (Burgess & Thompson, 2015), achieving sufficient power to confidently estimate the presence or absence of a causal relationship requires a large sample size; indeed, for most traits and outcomes, a minimum study size of tens of thousands of individuals is often required (Burgess, 2014; Freeman, Cowling, & Mary Schooling, 2013). The application of MR has therefore been limited to phenotypes with sufficiently large genetic datasets which happen to be available to the research community.

In this work, we apply contemporary techniques in machine learning to improve the power and sample-efficiency of MR by using individual-level genetic data. Specifically, we focus upon the identification and representation of instrumental variables by learning a valid and influential latent instrumental variable from candidate genetic variants. Our framework leverages recent work on learning and combining noisy labels (Natarajan, Dhillon, Ravikumar, & Tewari, 2018); (Ratner, De Sa, Wu, Selsam, & Ré, 2016; Ratner et al., 2018), with the goal of more accurate prediction and representation of population-level variance using individual-level data. Our approach, termed **I**nstrumental **V**ariable s**Y**nthesis (IVY), is complementary to existing causal effect estimators, whether classical (the Wald estimator (Wald, 1940), the two-stage least squares approach (Angrist et al., 1996)), robust (Bowden, Davey Smith, Haycock, & Burgess, 2016; Kang, Zhang, Cai, & Small, 2016), or modern (deep learning-based techniques (Hartford, Lewis, Leyton-Brown, & Taddy, 2017); IVY seeks to produce instrumental variables that can be used by downstream standard instrumental variable methodology for estimating causal-effects.

## 2. METHOD

Allele scores (Burgess, Dudbridge, & Thompson, 2016; Burgess & Thompson, 2013) are an effective approach to summarize the effects of multiple single nucleotide polymorphisms (SNPs) on polygenic traits such as lipids level and blood pressure. Given a set of SNPs {*w*_1_, *w*_2_, …, *w*_*p*_} as IVs, where *w*_*j*_ ∈ {−1,1} represents the absence or presence of an effective allele, allele score methods find a weight for each SNP and use the weighted combination of *w*_*j*_’s as the score value to describe the genetic contribution to the risk factor.

Conventionally, when allele scores are derived from the data, the risk factor *X* is used as a supervision signal to guide the derivation so that much of the variance in the risk factor can be explained by the allele score, increasing the power of the allele score. However, in observational studies, the value of the risk factor is confounded by many measured or unmeasured confounders. Therefore, when there are SNPs contributing to the allele score that are correlated with the confounder, we call such SNPs confounded SNPs, and they are invalid instrumental variables that should not be used to derive the allele score. Since the independence of a SNP from all potential confounders cannot be verified, conventional allele scoring approaches are particularly vulnerable to confounded SNPs. An allele score significantly corrupted by confounded SNPs could result in misleading estimate of causality.

### 2.1. Assumptions and Problem Formulation

In contrast to existing allele score approaches, IVY derives an allele score by circumventing the direct utilization of the risk factor as a supervision signal. Instead, IVY assumes that *w*_*j*_’s are noisy realizations of a hidden binary instrumental variable *z* ∈ {−1,1} that summarizes the genetic contribution to the observed risk factor. For each individual, IVY views that the status of *z* is reflected from the corresponding SNPs *w*_*j*_’s. When *w*_*j*_ = 1, *w*_*j*_ will consider that it is more likely that *z* = 1 and the genetic contribution to the observed risk factor is high. On the other hand, when *w*_*j*_ = −1, *w*_*j*_ will consider it is more likely that *z* = −1 and the genetic contribution to the observed risk factor is low.

The role of the hidden *z* modeled by IVY is comparable to the role of an allele score. This is because both *z* and the allele score can be viewed as summary variables of genetic risks. In this sense, deriving a hidden *z* in IVY from SNPs for summarization is similar in sprit to the goal of allele score methods. Furthermore, an allele score is also assumed to be a valid IV for the causal relationship in question. Since *z* is solely derived from the SNPs, when all the SNPs are valid IVs, the resultant *z* will also be a valid IV. This is also the case for allele scores.

IVY seeks to model the joint distribution of *w*_1_, *w*_2_, ⋯, *w*_*p*_, and *z*, without having access to *z*. Here we discuss the scenario where we assume that the SNPs are conditionally independent given *z*.

That is *w*_*j*_ ⊥ *w*_*j*_′ ∣ *z*, for all *j*, *j*′ ∈ {1,2, …, *p*}, and *j* ≠ *j*′. More specifically, for some parameters θ = (θ_*1z*_, θ_*2z*_, ⋯, θ_*pz*_, θ_*z*_), the joint distribution follows:

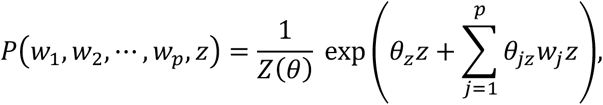

where *Z*(*θ*) is the partition function that ensures the appropriate normalization of the distribution.

It should be noted that the conditional independence assumption made in IVY is similar to the conditional independence assumption of *w*_*j*_’s given a binary risk factor in an ordinary allele score, as pointed out by (Sebastiani, Solovieff, & Sun, 2012). In practice, SNPs that are not in physical linkage are desirable for allele score methods. In IVY, we also made use of such SNPs. We refer interested readers to (Ratner et al., 2018) for a discussion of scenarios such as handling dependencies among *w*_*j*_’s. In this case, Ivy generalizes beyond the standard conditional independence assumption. Table 1 summarizes the comparison of modeling assumptions among Ivy, UAS, and WAS.

**Table 1.** Comparison of Modeling Assumptions among Ivy, UAS, and WAS

| Method | Dependencies of SNPs | Summary variable | Relationship between SNPs and the summary variable | Comments |
| --- | --- | --- | --- | --- |
| IVY | Conditional independence and beyond | Yes | Markov Random Field | Learns from individual-level data. Can be extended to handle dependencies among SNPs (Ratner et al., 2018). |
| UAS | Conditional Independence | Yes | Additive Model | IVY and WAS are similar to UAS when the effect of individual SNPs are of similar magnitude |
| WAS | Conditional independence | Yes | Additive Model | Can be learned from summary data. Conditional independence assumption (Sebastiani et al., 2012). |

Our parameterization of the joint distribution also implies the symmetry property that *P*(*w*_*j*_ = 1 | *z* = 1) = *P*(*w*_*j*_ = −1 | *z* = −1), for all *j* ∈ {1,2, …, *p*}. In practice, such an assumption can be roughly satisfied by excluding SNPs of low minor allele frequency.

### 2.2. Learning and Scoring

We now describe the learning algorithm of IVY and the method that IVY uses to synthesize an IV. Denote a random variable *a*_*j*_ = *w*_*j*_*z*. IVY first seeks to estimate 𝔼*a*_*j*_ without having access to *z*. Intuitively, *a*_*j*_ represents the agreement or disagreement between *w*_*j*_ and *z*. Therefore, the higher 𝔼*a*_*j*_ the more accurate is *w*_*j*_ in predicting *z*. Since *w*_*j*_′*s* are usually SNPs that associate with the risk factor and contribute to the overall genetic risk described by *z*, it is reasonable to view that *w*_*j*_ can predict *z* better than random guessing. This means 𝔼*a*_*j*_ > 0. One can further show that *a*_*j*_ and *a* ′ are independent, i.e. *a*_*j*_ ⊥ *a*_*j*_ ′ for all *j*, *j*^*′*^ ∈ {1,2, …, *p*}, where *j* ≠ *j′*. This means that Cov (*a*_*j*_, *a*_*j*_′)= 0. Furthermore,

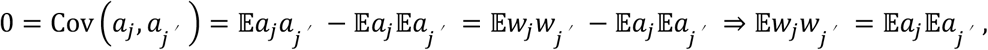

where we have used the fact that 𝔼*a*_*j*_*a*_*j*_ ′ = 𝔼(*w*_*j*_*z*) (*w*_*j*_′ *z*) = 𝔼*w*_*j*_*w*_*j*_′ *z*^2^ = 𝔼*w*_*j*_*w*_*j*_′.

Note that 𝔼*w*_*j*_*w*_*′*_ are the entries from the second moment matrix from *W*_*j*_’s, which are computable from *w*_*j*_’s alone. By assuming that all *w*_*j*_’s can predict *z* better than random guessing, that is, P(*w*_*j*_ = *z*) > P(*w*_*j*_ ≠ *z*) and hence

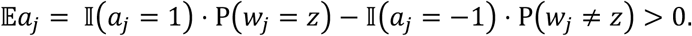

We can estimate log 𝔼*a*_*j*_’s by solving the following least square problem:

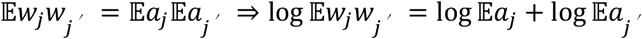

for all *j*, *j*^*′*^∈ {1,2, …, *p*}, and *j* ≠ *j′*, where we have used the fact that 𝔼*a*_*j*_ > 0 in order to take its logarithm.

Upon the estimation of log 𝔼*a*_*j*_, Ivy uses the following formula to compute the allele score:

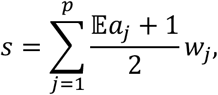

where 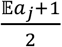 can be interpreted as the accuracy of *w*_*j*_ in predicting the value of z. In this way, IVY yields a synthesized IV that upweights SNPs that are more predictive of *z* while downweighting those that are less predictive. This is distinctive from conventional allele scores that tend to upweight SNPs that are more predictive of *x*.

## 2.3. IVY in Practice

In practical Mendelian randomization settings, 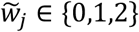, which represents the number of effective alleles. We use 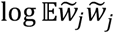’s in lieu of log 𝔼*w*_*j*_*w*_*j*_‣’s as data to construct the least square estimate of log 𝔼*a*_*j*_ and use the estimated accuracies to weight 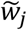 as an allele score.

In this way, IVY offers a data-driven solution to compute weights for SNPs to construct an allele score without the direct utilization of the confounded risk factor *x*. Since *z* is an IV, by definition, it is independent of a given confounder *c*. Even if we have a set of confounded SNPs {*v*_1_, *v*_2_, …, *v*_q_} that can be viewed as independent and noisy realizations of *c*, these confounded SNPs will be independent of {*w*_1_, *w*_2_, …, *w*_*p*_}, which is a set of valid instrumental variables that can be viewed as independent and noisy realizations of *z*. Therefore, when *q* ≪ *p*, the allele scores learned will not be compromised much by the confounded SNPs as most SNPs that are valid aim to learn the concept of *z*, and will down weight the few invalid SNPs that try to learn the concept *c* independent of *z*. Notably, these confounded SNPs can be highly dependent on the confounder, which in turn can highly influence the risk factor. Therefore, using those confounded SNPs can be very predictive of the risk factor and hence conventional allele scores will assign large weights on these invalid SNPs without scrutinizing their identities, leading to potentially erroneous causal estimates.

## 3. Experimental procedures using real-world data

We used data from the UK Biobank (UKB) as the basis of comparing IVY to standard methods for instrumental variable synthesis in MR. Our main experimental procedure is similar to 2-sample MR where candidate instrumental variables are derived from the summary statistics from separate external GWAS, and the causal effect estimate performed in a second independent GWAS. The European population of the UK Biobank (*n* ≈ 330,000) was randomly divided into two equally sized groups of individuals for 1. Re-estimation of weights for an instrumental variable and 2. Estimation of the causal effect (Figure 1). This procedure seeks to maintain the methodological rigor of 2-sample MR, whereby the cohort from which the instrumental variable weights are re-estimated and the cohort for estimation of the causal effect <u>remain completely separate</u>: only the re-weighted instrumental variable crosses between the two groups. While there is a proliferation of methods for estimating causal effects in MR, for simplicity we consider only causal estimates derived from a Wald-ratio as discussed above. We report the median and the median absolute deviation (MAD) associated with the Wald ratio estimate. The median and MAD are generated by repeating the aforementioned random division of the dataset as well as the instrumental variable reweighting and causal effect estimation procedure 1000 times.

**Figure 1:**
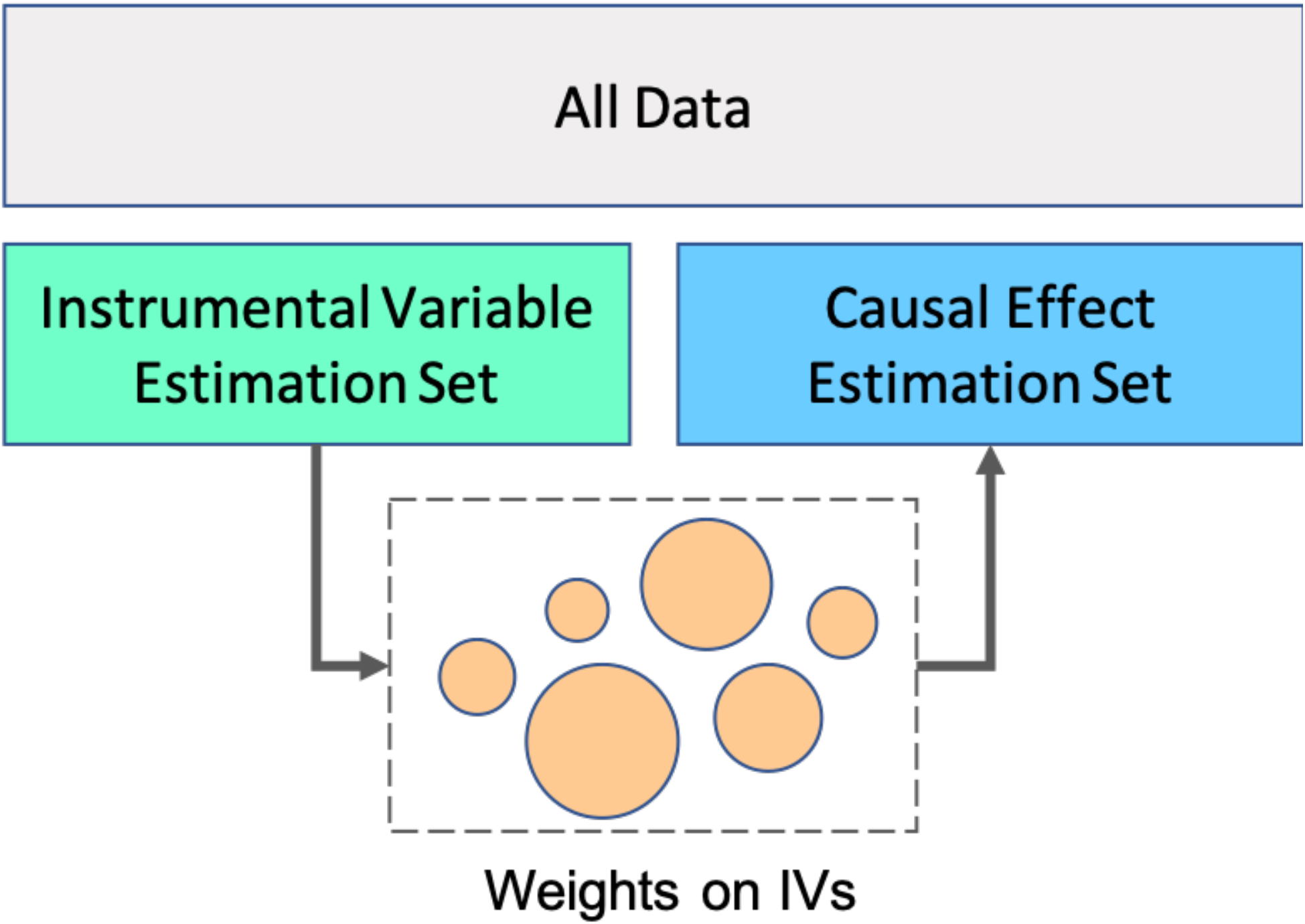
Experimental procedure using the UK Biobank for evaluation of IVY in the format of two-sample Mendelian Randomization.

To better understand the performance of IVY relative to standard methods we conducted sub-sampling experiments. Random sub-samples were taken from the instrumental variable estimation set to evaluate the robustness of the re-weighted instrument for capturing a proportion of the variance. Random sub-samples were taken from the causal effect estimation set to evaluate the relative power of the re-weighted instrument for correct and accurate estimation of a causal effect or absence thereof.

To test the performance of IVY on empirically collected data, we analyzed the causal relationship between systolic blood pressure (SBP) and coronary artery disease (CAD) using Mendelian randomization. CAD is a well characterized outcome with well-described causal and non-causal relationships, and previous Mendelian randomization analyses have provided strong support for a well-established causal relationship between blood pressure and risk of CAD (Bowden et al., 2016). Briefly, twenty-six candidate variants were selected from external GWAS for systolic blood pressure (International Consortium for Blood Pressure Genome-Wide Association Studies et al., 2011; Lieb et al., 2013; Wain et al., 2011), to be used as instruments for risk factors (instrumental variables related to SBP). Those 26 uncorrelated variants showed genetic associations with SBP in over 200,000 individuals at genome-wide significance (p<5e-8) (International Consortium for Blood Pressure Genome-Wide Association Studies et al., 2011) and have been further validated on data from the CARDIoGRAM consortium (Lieb et al., 2013).

To rule out the possibility that IVY can fit artifactual causal associations (false positives), a negative control experiment was conducted. Mendelian randomization procedures using weighted and unweighted allele scores were compared to IVY on the task of determining the causal relationship between SBP and an unrelated outcome, where no causal effects are expected. Specifically, CAD was replaced by a genetically unrelated phenotype (as determined by LD score regression (Bulik-Sullivan et al., 2015)) and the presumably null causal relationship between SBP and this phenotype was tested using the 26 SBP variants to build instrumental variables. Bone disease was chosen as negative control outcome based on the analysis of hundreds of phenotypes through the LD Hub (Zheng et al., 2017) (LD score regression correlation with CAD: r=0.0009, S.E.=0.151, p=0.995).

We further investigate the utility of IVY in correctly detecting non-causal correlations. As an example, we consider the strong observational association between high density lipoprotein (HDL) and CAD. In observational data, increased HDL appears to be protective from CAD, and the strong negative association between HDL and CAD would suggest that raising HDL level can have protective effect against the occurrence of CAD. However, such an association is spurious, as demonstrated by various clinical trials and Mendelian randomization studies (Schwartz et al., 2012; Voight et al., 2012). We seek to replicate the dismissal of this spurious correlation via IVY by making use of the 19 SNPs reported in (Holmes et al., 2015) to synthesize the instrumental variables.

Finally, we study the performance of IVY under relatively agnostic criteria for inclusion of SNPs reported in large-scale GWAS. We examine the estimates of causal effect provided by IVY across a range of associated variants from the GWAS catalog. To this end, we consider the SBP→CAD relationship, where we make use of 238 SNPs associated with SBP as reported in the GWAS Catalog (Buniello et al., 2019), a knowledge base that curate SNPs associated with various biomarkers from published GWAS. We use subsets of increasing number of SNPs among the 238 SNPs to synthesize instrumental variables and notice the decreases in the uncertainty of the resultant causal effect estimates. We further consider ascertaining the relationships between four serum biomarkers (total cholesterol, C-reactive protein, triglycerides, and vitamin-D) and CAD using SNPs associated with these four risk factors respectively. Given a study population of European ancestry in the UK Biobank, only SNPs arising from studies which included individuals of European ancestry were included.

## 4. RESULTS

We sought to compare the performance of IVY in identifying causal relationships between SBP and CAD with the performance of standard methods UAS and WAS using different number of samples in the instrumental variable estimation set and causal effect estimation set (Figure 2). As a negative control, we estimate a putatively non-existent causal relationship between SBP and bone density disorder (BDS, Figure 3). We investigate whether IVY can dismiss spurious correlation using the confounded spurious relationship between high-density lipoprotein (HDL) and CAD as an example (Figure 3). We then show that IVY can deliver better powered analysis by incorporating more associated SNPs to the risk factor in Table 2. Finally, we conduct causal effect estimation using un-curated SNPs for the relationships between four serum biomarker and CAD (Table 3).

**Table 2:** Variance decreases but causal effect estimates remain stable across increasing subsets of SNPs included by IVY.

| # SBP SNPs (fraction total) | Median Causal estimate | Median absolute deviation |
| --- | --- | --- |
| 15 (1/16) | 0.90 | 0.38 |
| 30 (1/8) | 0.90 | 0.27 |
| 60 (1/4) | 0.92 | 0.17 |
| 119 (1/2) | 0.91 | 0.10 |

instrumental variable estimation set and causal effect estimation set sizes each 166,499.

**Table 3:** IVY correctly ascertains causal relationships between four serum biomarker and coronary artery disease.

| Serum biomarker | # SNPs in GWAS Catalog | Median Causal estimate | 95% Confidence Interval | Ground truth |
| --- | --- | --- | --- | --- |
| Total cholesterol | 279 | 0.851 | [0.623, 1.077] | Increased risk |
| C-reactive protein | 88 | 0.176 | [-0.257, 0.594] | Non-causal |
| Triglycerides | 273 | 1.777 | [1.520, 2.103] | Increased risk |
| Vitamin-D | 25 | 0.157 | [-0.182,0.505] | Non-causal |

CE and instrumental variable cohort sizes each 166,499.

**Figure 2:**
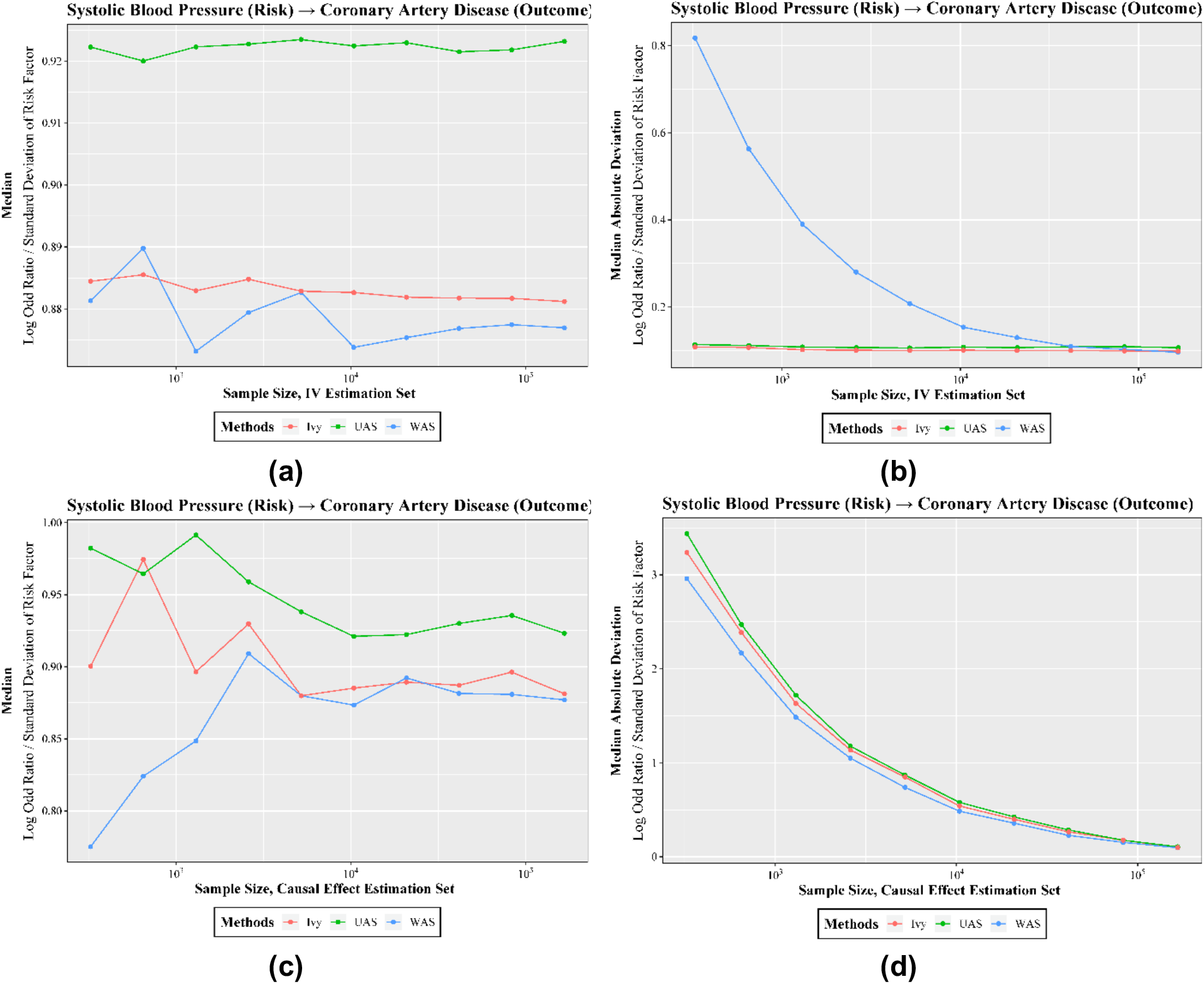
IVY correctly determines a causal effect between systolic blood pressure and coronary artery disease at different sample sizes.

**Figure 3:**
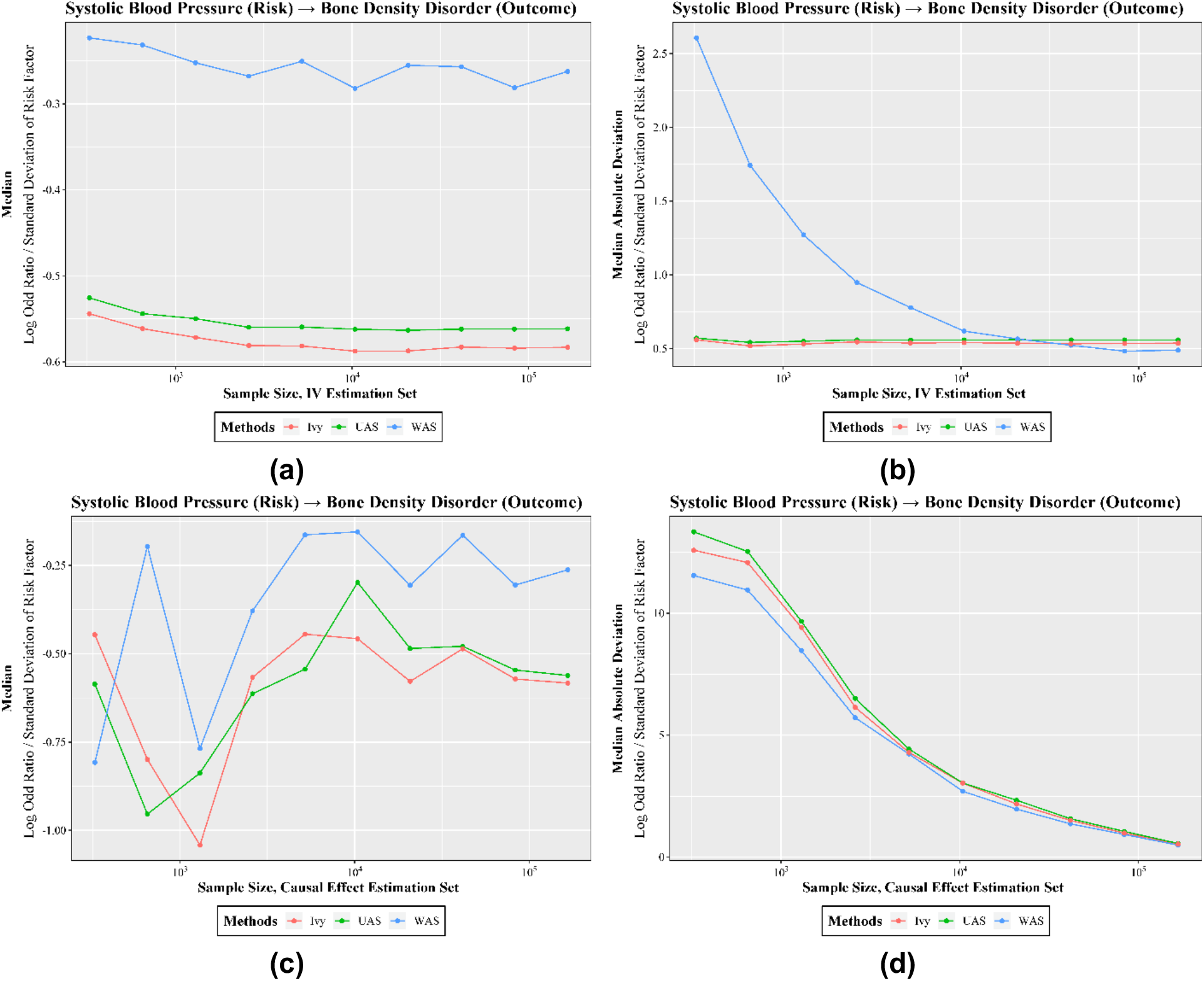
IVY correctly determines the absence of a causal effect between systolic blood pressure and bone disease at different sample sizes.

### 4.1. Comparing IVY with UAS and WAS for known causal relationships

We report our comparison between IVY and allele scores for known causal relationships using SBP→CAD as an example. IVY faithfully recapitulates known causal relationships between SBP and CAD similar to both UAS and WAS. When the size of the instrumental variable estimation dataset is varied, causal estimates from the WAS become less precise (indicated by larger MAD) with less data but are invariant when using instrumental variables derived from IVY and UAS of different sizes. This is reflected in (Figure 2b), where as the sample size decreases, the MAD of WAS increases substantially, while the MAD of UAS and IVY maintains relatively stable. Functionally, so long as the alleles are present in the instrumental variable estimation set no re-weighting of the UAS occurs. When the size of the CE dataset is varied, IVY provides accurate estimates of the SBP→CAD causal relationship similar in magnitude to the WAS which are lower than the UAS (Figure 2d). The causal estimates from all 3 methods remain stable to below 10^4 individuals but precision decreases with sample sizes below 10^5 individuals (Figure 2b and Figure 2c).

To demonstrate that IVY does not capture artificial causal associations (false positives), we consider the relationship between SBP and BDS as an example. IVY correctly detects an absence of causality between blood pressure and bone disorders similar to UAS and WAS. The correct estimate for an absence of effect becomes more precise in larger instrumental variable estimation cohort sizes using all three methods (Figure 3). Overall, IVY may provide a method of estimating instrumental variables which provides a more stable estimate of CE than WAS when smaller sample sizes are provided for re-weighting the instrumental variable and is more robust to inflation of the CE than the UAS. IVY retains the both the advantages of UAS and WAS while dispensing with the disadvantages.

### 4.2. Comparing IVY with UAS and WAS for a known non-causal relationship

To test the performance of IVY in a known non-causal relationship we examined HDL and CAD. In our experimental protocol, IVY can successfully detect the true absence of a causal relationship between HDL and CAD (Figure 4) despite the strong observational association. Although UAS and WAS can also detect the absence of a causal relationship, IVY yields a median causal effect that is closer to the null and when ablating either the IV estimation set (Figure 4a) or the causal effect estimation set (Figure 4c). The variance of the estimate as indicated by the MAD (Figure 4b and Figure 4d) of the three methods are similar to that of the previously described causal and non-causal relationships (Figures 2,3) where the WAS appears to be most sensitive to the sample size of the IV estimation set while the three methods display similar characteristics when varying the size of the causal effect estimation set.

**Figure 4:**
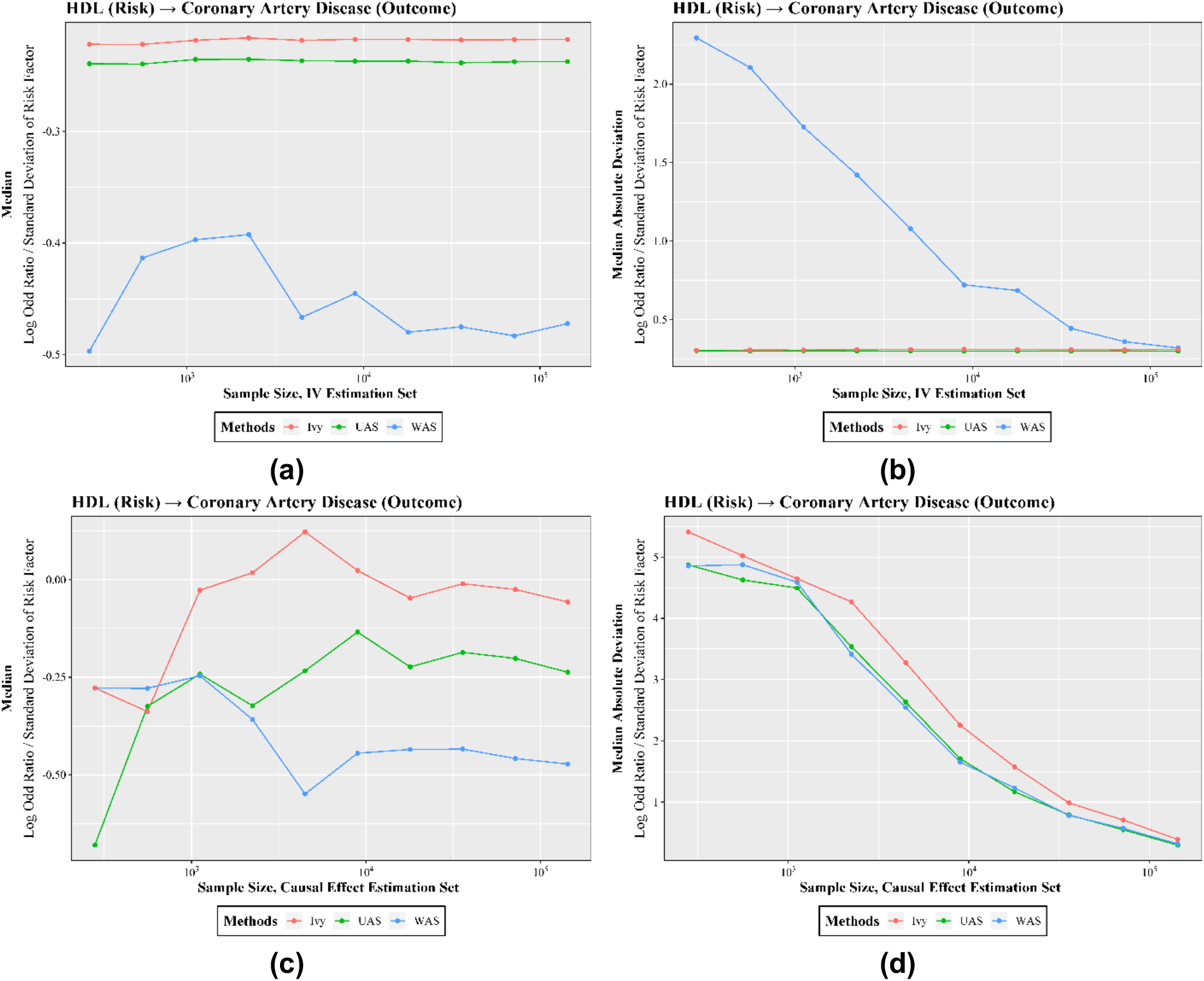
IVY correctly determines the absence of a causal effect between HDL-cholesterol and coronary artery disease at different sample sizes.

### 4.3. Incorporating more associated SNPs may yield better powered study with IVY

When performing two-sample MR, a greater number of SNPs independently associated with risk factor increases the proportion of variance explained and improves the certainty of the causal estimate. In order to examine the performance of IVY utilizing different subsets of SNPs, we measured the causal estimate and median absolute deviation (MAD) across subsets of SNPs of decreasing size for a single trait. We consider the SBP→CAD relationship, for which we find 238 SNPs associated with SBP in the GWAS catalog as candidate instrumental variables. We carry out causal effect estimate over subsets of the 238 SNPs. Specifically, we fix the split between the instrumental variable estimation set and the causal effect estimation set. Within the instrumental variable estimation set, we randomly select 1/16, 1/8, 1/4, and 1/2 of all the 238 SNPs respectively to derive the IV. We repeat this random selection procedure for 1000 times in order to compute median and MAD. We expect that a larger subset of SNPs corresponds to lower MAD (higher certainty) of the estimate. The experimental results are summarized in Table 2, where we observe that as the number of instrumental variables used increases, the causal effect remains relatively constant while the MAD decreases. These results suggest that the causal estimates derived from the IVY procedure behave in an expected manner; increasing the number of SNPs within the instrumental variable explains more variance and delivers a better powered instrumental variable for MR when using the IVY framework.

### 4.4. Ascertain relationships between four serum biomarkers and CAD using un-curated SNPs via IVY

Given the IVY procedure is sensitive to the number of included SNPs and proportion of variance explained in the risk factor, we investigate whether IVY is capable of estimating causal effects using SNPs with little or no curation of candidate instrumental variables. To this end, we consider the known relationships between four additional serum biomarkers and coronary artery disease for which the ground truth is known. We use the GWAS catalog to identify all SNPs that are associated with each of the four serum biomarkers. Including all the available SNPs in the GWAS catalog, we estimate the causal effects for each of these relationships using IVY. The experimental results are summarized in Table 3, where IVY is capable of estimating causal effects in a manner consistent with the ground truth when including un-curated SNPs.

## 5. DISCUSSION

The method proposed here IVY provides a framework for instrumental variable synthesis for causal inference that leverages weak supervision to learn a hidden instrumental variable from the vastly available candidate instrumental variables of potentially lower quality in the data.

Through empirical demonstrations we demonstrate that IVY can recover causal effects in a manner that retains the desirable characteristics of conventional weighted and unweighted allele scoring for instrumental variables in a framework analogous to two-sample Mendelian randomization. Through ablating the instrumental variable estimation set and causal effect estimation set, we show that IVY has the potential to deliver overall better powered MR using individual-level data in a more sample-efficient fashion.

The underlying statistical assumptions made in IVY are connected to the assumptions made in conventional allele scores. Specifically, the conditional independence assumption made in IVY for the inclusion of individual SNPs is valid because the same assumption is made for methods employing allele scores (Sebastiani et al., 2012). We point out the possibility of extending beyond the assumption of conditional independence such as handling SNPs occurring physically upon the same chromosome which display some degree of genetic linkage (Ratner et al., 2018). Empirical validation of such an extension is left for future research.

We offer intuitive explanation on why IVY can potentially synthesize an instrumental variable that is more robust to invalid SNPs. Specifically, as the re-weighting procedure performed by IVY is not supervised by a potentially confounded risk factor, IVY may avoid over-weighting an invalid confounded SNPs during the synthesis. Our findings illustrate that when smaller sample sizes are provided for re-weighting of the instrumental variable, IVY provides a more stable estimate of the CE than the WAS method. Additionally, the instrumental variable produced by IVY appears to be more robust to inflation of the CE than the UAS method. In this manner when the Wald ratio is used to estimate the causal effects, the CE produced by IVY appear to be both more stable and resistant to inflation compared to standard methods. Additional research is necessary to determine appropriate procedures for including the re-weighted instrumental variable provided by IVY with weighted median and other accepted methods of computing CE estimates (Bowden et al., 2016).

The IVY framework as described here is not without limitations. To begin with, the assumption of the existence of a hidden binary valid instrumental variable *z* is somewhat different than conventional allele scores that seek to synthesize a numeric score as an instrumental variable. Nonetheless, the potential limitation of this assumption can be mitigated by considering the log probability of *z* = 1, i.e. log *P*(*z* = 1|*w*_1_, *w*_2_, ⋯, *w*_*p*_), which is a numeric quantity, as an instrumental variable to be synthesized. As a result, the scoring formula that gives *s* as a weighted combination of estimated accuracies of *w*_*j*_’s proposed in the paper can also be replaced by this log probability. Secondly, our investigation assumes the existence of only one hidden instrumental variable. While such an assumption is consistent with that of allele scores due to the similar conditional independence assumption between the two, it is desirable to generalize the IVY framework where multiple hidden instrumental variables are available.

Finally, the analysis conducted in this paper is only among population of European ancestry. Further investigation is needed to ensure that the framework of instrumental variable synthesis presented here is not derive power from artifacts of European population substructure and is generalizable to individual-level data of other ancestries.

Overall, given the stability offered by our proposed methods with relatively small sample sizes for re-weighting of the instrumental variable or estimating a causal effect, we envision that IVY may be useful to perform MR in small population sizes. Real world examples of where it may be desirable to perform MR in small population sizes might include cultural, ethnic, and geographic groups not represented in large GWAS studies, sub-analyses of larger datasets stratified by sex or a specific clinical risk factor, or for risk-factors or outcomes which have been described in relatively small GWAS. An R-package for this procedure is forthcoming.

## 6. Appendix

**Figure 5:**
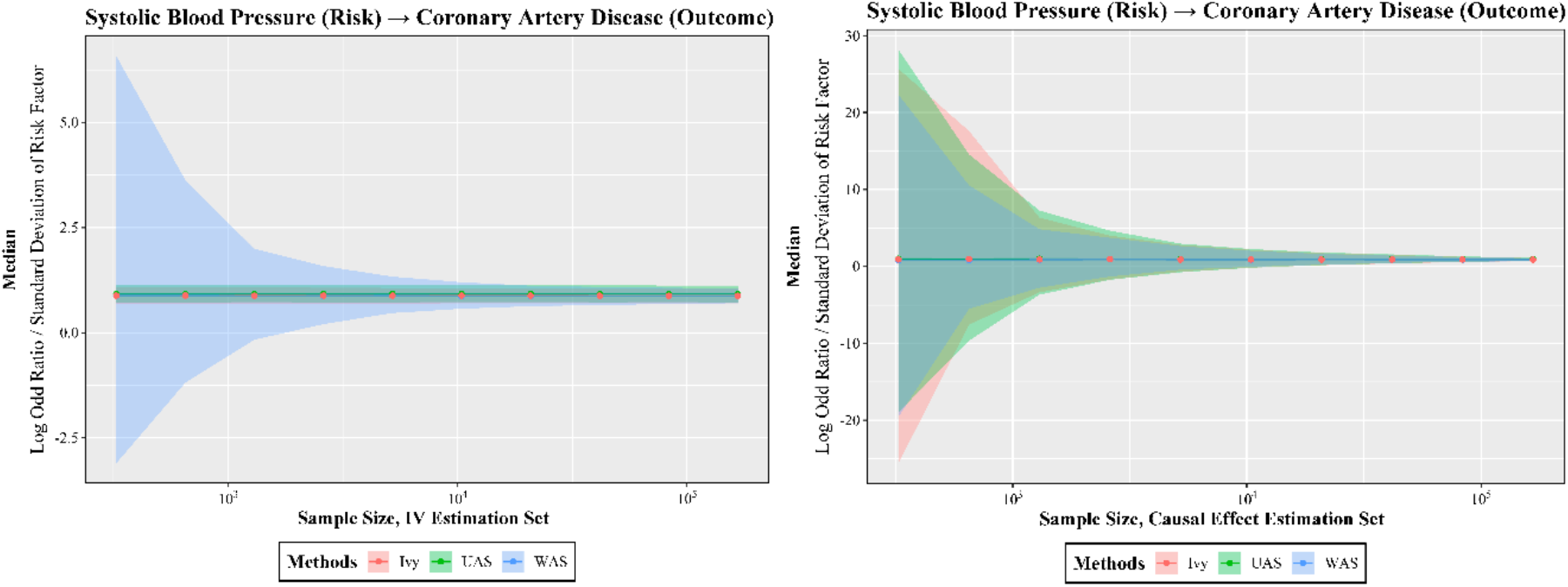
Ribbon Plots Corresponding to Figure 2. IVY correctly determines a causal effect between systolic blood pressure and coronary artery disease at different sample sizes.

**Figure 6:**
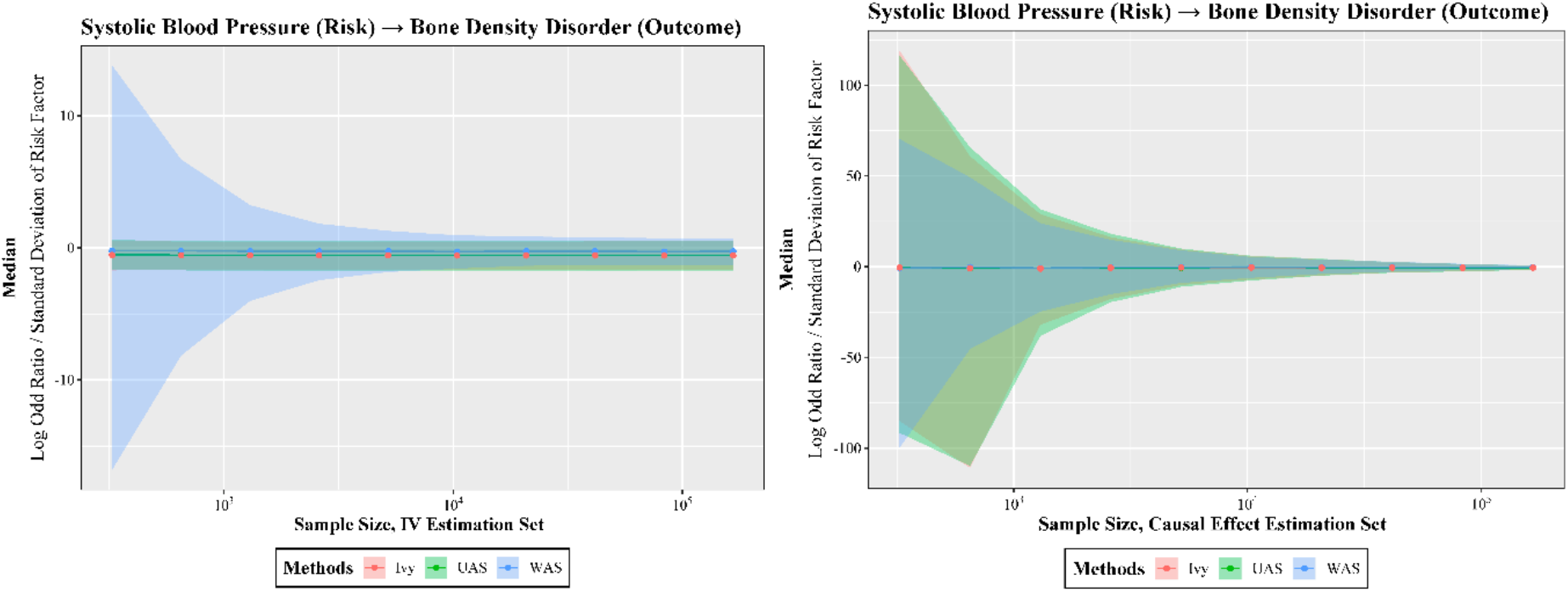
Ribbon Plots Corresponding to Figure 3. IVY correctly determines the absence of a causal effect between systolic blood pressure and bone disease at different sample sizes.

**Figure 7:**
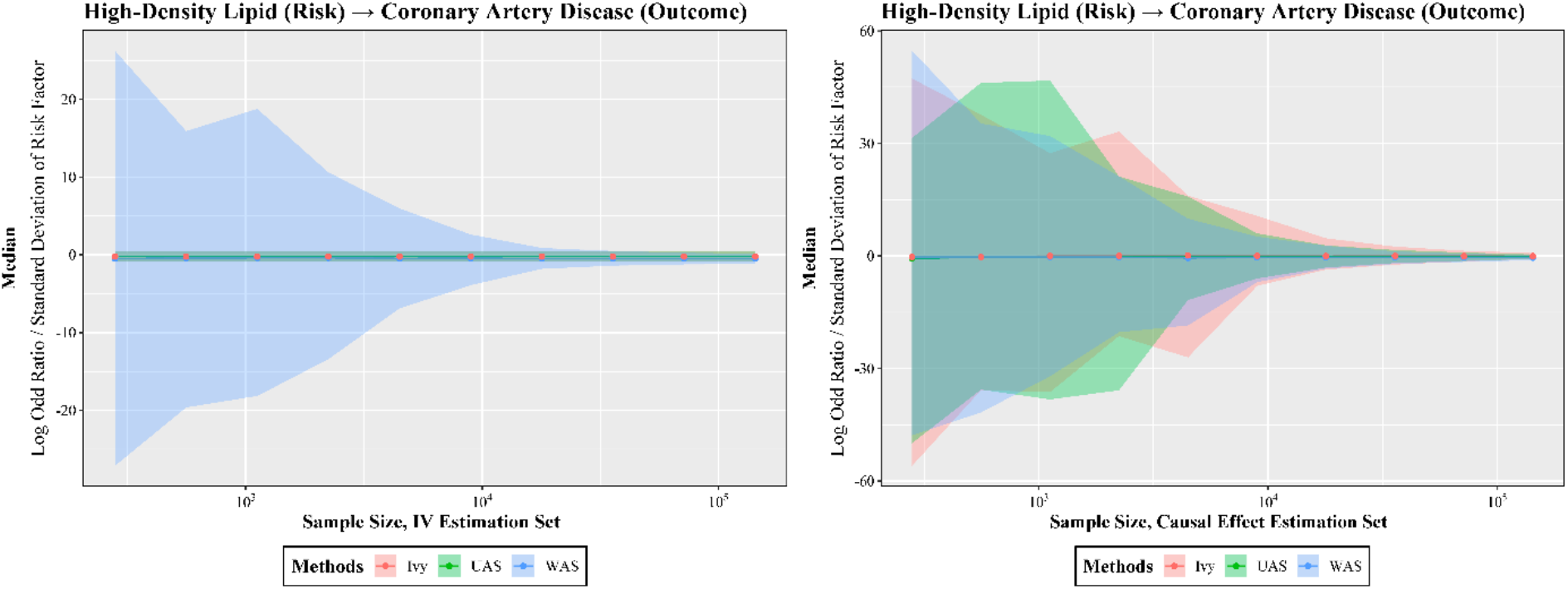
Ribbon Plots Corresponding to Figure 4. IVY correctly determines the absence of a causal effect between HDL-cholesterol and coronary artery disease at different sample sizes.

